# bpRNA-align: Improved RNA Secondary Structure Global Alignment for Comparing and Clustering RNA Structures

**DOI:** 10.1101/2022.03.23.485538

**Authors:** Brittany Lasher, David A. Hendrix

**Affiliations:** Department of Biochemistry and Biophysics, Oregon State University; School of Electrical Engineering and Computer Science, Oregon State University

## Abstract

Ribonucleic acid (RNA) is a polymeric molecule that is fundamental to biological processes, with structure being more highly conserved than primary sequence and often key to its function. Advances in RNA structure characterization have resulted in an increase in the number of accurate secondary structures. The task of uncovering common RNA structural motifs with a collective function through structural comparison remains challenging and could be used to discover new RNA families. In this work, we present a novel secondary structure alignment and clustering method, bpRNA-align. bpRNA-align is a customized global structural alignment method, utilizing an inverted (gap extend costs more than gap open) context-specific affine gap penalty and a structural, feature-specific substitution matrix to provide similarity scores. We evaluate our similarity scores in comparison to other methods, using affinity propagation clustering, applied to a benchmarking data set of known structure types.

## Introduction

Ribonucleic acid (RNA) consists of polymeric chains of nucleotides that form structures through base pair interactions. RNA has numerous biological roles including encoding information, as in messenger RNAs (mRNAs), decoding information (e.g. transfer RNAs, tRNAs), and regulation of gene expression. Noncoding RNAs (ncRNAs) are RNAs that, unlike mRNAs, do not encode proteins. ncRNA structure is more highly conserved than primary sequence and defines their function. With advances in RNA structure characterization through X-ray crystallography [1], cryogenic electronic microscopy [2], nuclear magnetic resonance (NMR) [3], and secondary structure (SS) predictions guided by RNA probing data, an increased quantity of accurate SSs have become available. In the Protein Data Bank (PDB) alone, over 50% of the RNA-only structures have been identified since 2010 [4]. The past decade has also seen the rise of high-throughput structure probing assays that have generated experimentally supported structures for over 150 unique transcriptomes [5].

The rising tide of available RNA structure data necessitates the need for improved approaches to compare and identify similar structures, and to organize available data. A popular strategy to compare RNA structures is through tree- or forest-based alignments, which are computationally expensive. RNAforester, is a comparison method which applies a forest-based alignment approach to simultaneously align structure and sequence [6,7]. Under this method, a SS is represented as a rooted ordered forest, which contains “P” nodes representing the base paired bonds, with sequence in the leaves. The RNAforester global structure alignment has made vast improvements to its algorithm, resulting in a time complexity of 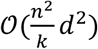, where *n* is the number of nodes in the forest, and *d* is the degree of the forest and *k* splits a forest at all possible points [8]. This approach can be challenging to understand and lacks distinct representations of unpaired loop types at the nucleotide-level (e.g. hairpin loops vs multiloops).

BEAGLE [9], another structural comparison method, utilizes the BEAR encoding to represent structural information as blocks characterizing different SS types (loop, internal loop, stem, or right/left bulge) and their lengths [10]. For example, stems of length 5 are represented as S_5_, and are encoded into the BEAR representation using a corresponding letter of the alphabet. This results in an overly complex representation, involving a different letter of the alphabet for each combination of length and structure type, where a limited maximum length is fixed for each structure block. Although, structural direction differentiation is implemented for bulges, with right and left bulge options, this is not applied for stem structure blocks, which could lead to misalignments as structures become more complex. BEAGLE aligns its structure representation using a modified Needleman-Wunsch sequence alignment algorithm, with generation of a log-odds substitution matrix for scoring alignments [9,10]. The large alphabet within the BEAR encoding requires BEAGLE to have a large substitution matrix with many terms. The challenge remains to develop fast and accurate SS comparison algorithms that still capture fine detail in their structure representations.

In our prior work we developed an automated structure annotation method, bpRNA, to label the structural features of RNA, including multiloops, bulges, hairpin loops, internal loops, and external loops, and pseudoknots [11]. bpRNA outputs an RNA SS representation with single-character structure codes for each nucleotide, referred to as a “structure array” (Fig 1A). The structure array provides a representation with several advantages compared to other methods. First, it uses a much smaller alphabet with single-character code corresponding to each position of the sequence. In other words, a four-nucleotide hairpin loop would be represented as HHHH, rather than using a different character for loops of different lengths [10]. Second, it explicitly incorporates multiloops, external loops, and ends into the sequence, rather than using the same character for each. Third, the structure array integrates well with dot-bracket representations because they are the same length. Within this work we utilize this structure array as an input to globally align RNA structures.

**Fig 1:**
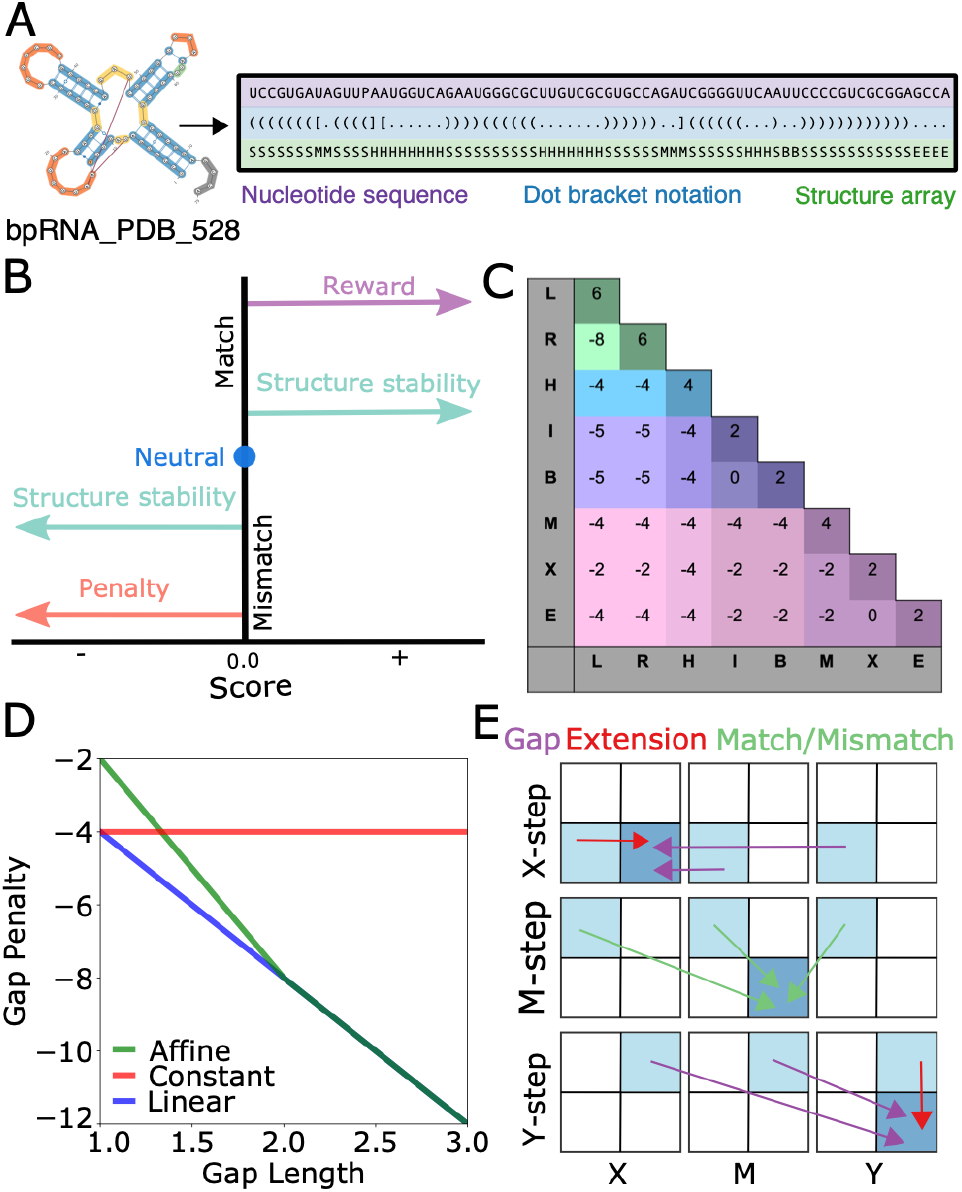
A. Needleman Wunsch algorithm customizations. Primary sequence (purple), dot bracket structure notation (blue) and bpRNA array structure representation (green) for the structure bpRNA_PDB_528. **B**. Scoring strategy used to generate the substitution matrix for the customized NW alignment approach, with positive scores correlated with their contribution to the structure stability, and with missmatch scores negatively correlated with their structure stability. **C**. Phenomenalogical scoring matrix generated from the scoring strategy. Colors and shades represent similar structural feature types. **D**. Approaches to handle gaps in the alignment strategy. **E**. Alignment moves between matrices for gaps, extensions, and match and mismatches.

We present bpRNA-align, a fast, easy-to-use program that identifies global similarity between RNA structures. bpRNA-align addresses previous RNA comparison limitations by using the bpRNA structure array, the length of which is the same as the RNA, and uses a small alphabet to represent all structure types. While affine gap penalties normally apply a greater penalty for gap openings than gap extensions, we do the reverse because single-nucleotide insertions or deletions are less destabilizing to RNA structure (e.g. a 1-nt bulge). Furthermore, we have also implemented context-specific gap penalties into bpRNA-align, which allow for natural variability observed within RNA structural features, enabling accurate identification of globally similar RNA structures, capable of connecting function to structure.

## Materials and Methods

### Alignment Algorithm

The bpRNA structural annotation tool identifies structural features, and creates the structural array sequence that includes stems (S), internal loops (I), bulges (B), hairpins (H), multiloops (M), external loops (X), and ends (E). For our bpRNA-align global alignment method, we further enhanced the bpRNA structure arrays using the dot-bracket sequence to covert the stem characters into left- and right-handed stem characters, L, and R. Our approach is to align structure arrays, uses Needleman-Wunsch alignment algorithm [12] as a basis for our global structure alignment approach. This is an application of dynamic programing, where a larger task is broken down into smaller sub-tasks, which are each solved to obtain the overall result. The Needleman Wunsch algorithm will align sequences *x* and *y*, which involves recursively computing a score matrix and a traceback matrix indicating the direction of each recursion step. Tracing backwards through this matrix, based on the directional pathway starting from the lower right position, provides the optimal alignment of *x* to *y*.

### Substitution matrix

To identify the scores assigned for match and mismatch results in the Needleman Wunsch alignment algorithm, a substitution matrix, *s*(*a, b*), is required, where *a* and *b* are the possible characters within the sequences *x* and *y*. Since a substitution matrix consisting of structural features as characters did not currently exist for scoring RNA structure arrays, it was necessary for us to develop one. We accomplished this through a phenomenological approach based on domain-specific knowledge and physical intuition about RNA structure. To assign matched and mismatched scores to the matrix we followed a reward and penalty strategy (Fig 1B). High rewards were applied for matching elements that were key to the structural integrity, and high penalties for mismatches which would significantly disrupt the structure. For example, matching right-sided stem characters would have a high reward. Conversely, mismatching a right-sided stem with a left-sided stem would result in a high penalty. In cases of mismatches of structural characters that are similar in nature, such as I and B, we applied a small or neutral penalty. The resulting substitution matrix is shown in Fig 1C using terms color-coded by similar structure types.

### Affine gap penalty

The affine gap penalty is a strategy commonly used in sequence alignment methods [13,14], which allows for penalizing gap openings with a parameter *G* < 0 different from gap extensions with parameter *E* < 0 (Fig 1D). Implementing affine gaps into dynamic programming alignment involves keeping track of 3 individual matrices, one for each type of step in the recursion (Fig 1E). Computing the optimal alignment includes diagonal steps along the *M*-matrix that include match or mismatch scores, *s*(*x*_*i*_, *y*_*j*_), horizontal steps along the *X*-matrix that correspond to gaps in *x*, and vertical moves along the *Y*-matrix that correspond to gaps in *y* (Fig 1E). Equations 1-3 show the system of recursion relations used for each possible step within the *M, X*, and *Y* matrices; where *i*, and *j* represent the position along the sequences *x* and *y*, respectively.

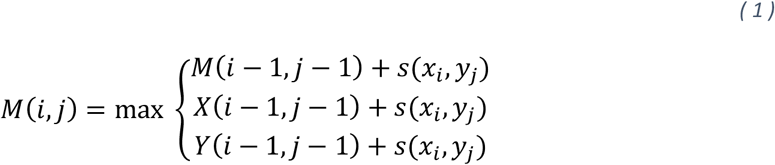

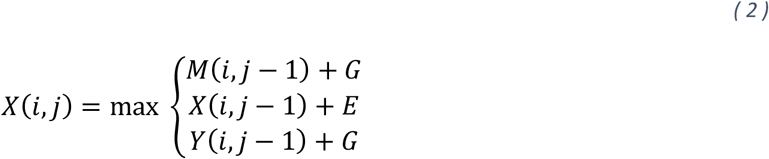

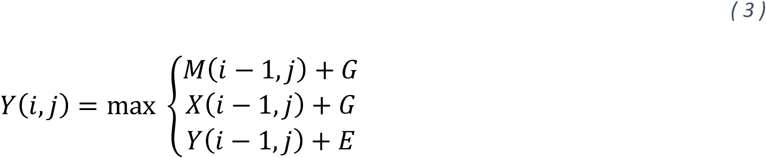

We implemented the affine gap penalty [13] in our method to allow for small gaps intermittently within the structure while still penalizing long stretches of gaps. This was achieved by setting a small gap opening penalty and a large gap extension penalty. This is opposite of traditional sequence alignment methods, which tend to assign a greater penalty to the insertion of a gap compared to extension of a gap [15].

### Natural variability within structural feature types and Context-specific gap penalties

To determine the variability within each structural feature type, we analyzed 26 different RNA families within the bpRNA-1m meta-database [11]. For each family the most common feature pattern was determined within the data. A feature pattern represents the feature and the feature-number in the order in which they appear in the structure but is not feature length specific. Thus, a structure composed two stems on either side of a hairpin loop, would be represented as S1H1S1. All unique structures in the family having the common feature pattern were used to determine the mean absolute deviation (MAD) of the length of each instance of a feature (e.g. H1, H2, H3, E1, E2, S1, S2, S3). We quantified the natural variability within each structure feature (S, H, I, M, E, X, B) using by computing the mean MAD over all the RNA families.

To compute context-specific gap penalties, we defined gap extension penalty *E* based on the mean MAD values for each structural feature, and the gap open penalty *G* to be *E*/2. To determine gap penalties within an adequate range and weighted based on their variability defined by their mean MAD value, we constructed a linear range of gap extension penalties between *E*_*min*_ = −1 and *E*_*max*_ = −7. These upper and lower bounds were chosen to be of a comparable weight to the scores in the substitution matrix. The trend used to determine the gap extension penalties is shown by the linear equation

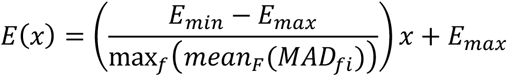

The *x* values represent the mean MAD value of a specific feature *f*, and *E*(*x*) values are the resulting gap extension penalties for the corresponding feature. In this equation, we first take the MAD length of each feature instance *fi* (e.g. H1, H2, S1), and then take the mean over all RNA families *F*. The denominator has the maximum *mean*_*F*_(*MAD*_*fi*_) length over all feature types *f*. Therefore, when *x* is zero, *E*(*x*) = *E*_*max*_, and when *x* is max_*f*_(*mean*_*F*_(*MAD*_*fi*_)), then *E*(*x*) = *E*_*min*_. Within our algorithm, we apply these context specific gap penalties for a particular feature type when the positions in each structure *i* and *j* are are inside, but not bordering, the same feature type region.

### Banded alignment

When structure arrays *x* and *y* have similarity to each other, we expect the optimal alignment path to be roughly diagonal. In these cases, the banded alignment strategy can be applied, minimizing the search space to a band around the diagonal of (half) width *w*. In our global alignment approach, we make use of the banded alignment strategy to increase speed. First, we select the longer of the two structure arrays, defining *n* = max (*length*(*x*), *length*(*y*)). The value of *w* is set to be *w* = 25 for our benchmark clustering tests. In practice, too narrow of a *w* could limit accuracy, but may also be more selective for highly similar structures. In practice, the *w* is specified by the user, and *w* ≈ *n*/4 sets a wide enough band to account for length variability.

### Benchmark data sets

We generated 4 different data sets, containing a range of RNA families or functional categories within bpRNA-1m [11], to test the performance of bpRNA-align. For each category in each data set up to 8 (dependent on the number of available structures) unique structures were randomly selected. The first data set generated was composed of 6 different RNA riboswitch categories, including Glycine, PreQ1-II, Purine, drz agam-1, SMK Box, and SAM (Fig 2A). The second dataset is composed of microRNAs including MIR159, mir-544, mir-1937, mir-166, mir-286, mir-720, and mir-684 (Fig 3A). These 8 different categories were composed of 7 unique structures for each category, all of which consist of a single segment. For the 3^rd^ and 4^th^ datasets, RNA families were chosen based on a consistent average number of structural segments, where a structural segment was previously defined [11] as a collection of adjacent or near-adjacent base pairs, only being interrupted by bulges, hairpin loops, or internal loops, but not multiloops. Dataset 3 consists of three-segment structures, including HCV_X3, SCARNA16, SNORA22, His_leader, 5S, PyrR, and ROSE (Supplementary Figure 3A). For each family in this dataset, 8 unique structures were selected. Dataset 4 consisted of structures with four segments, including radC, tfoR, MicX, RsaD, BTE, and rpsL_psuedo (Supplementary Figure 4A). For each family in dataset 4, 7 unique RNA structures were included.

**Fig 2:**
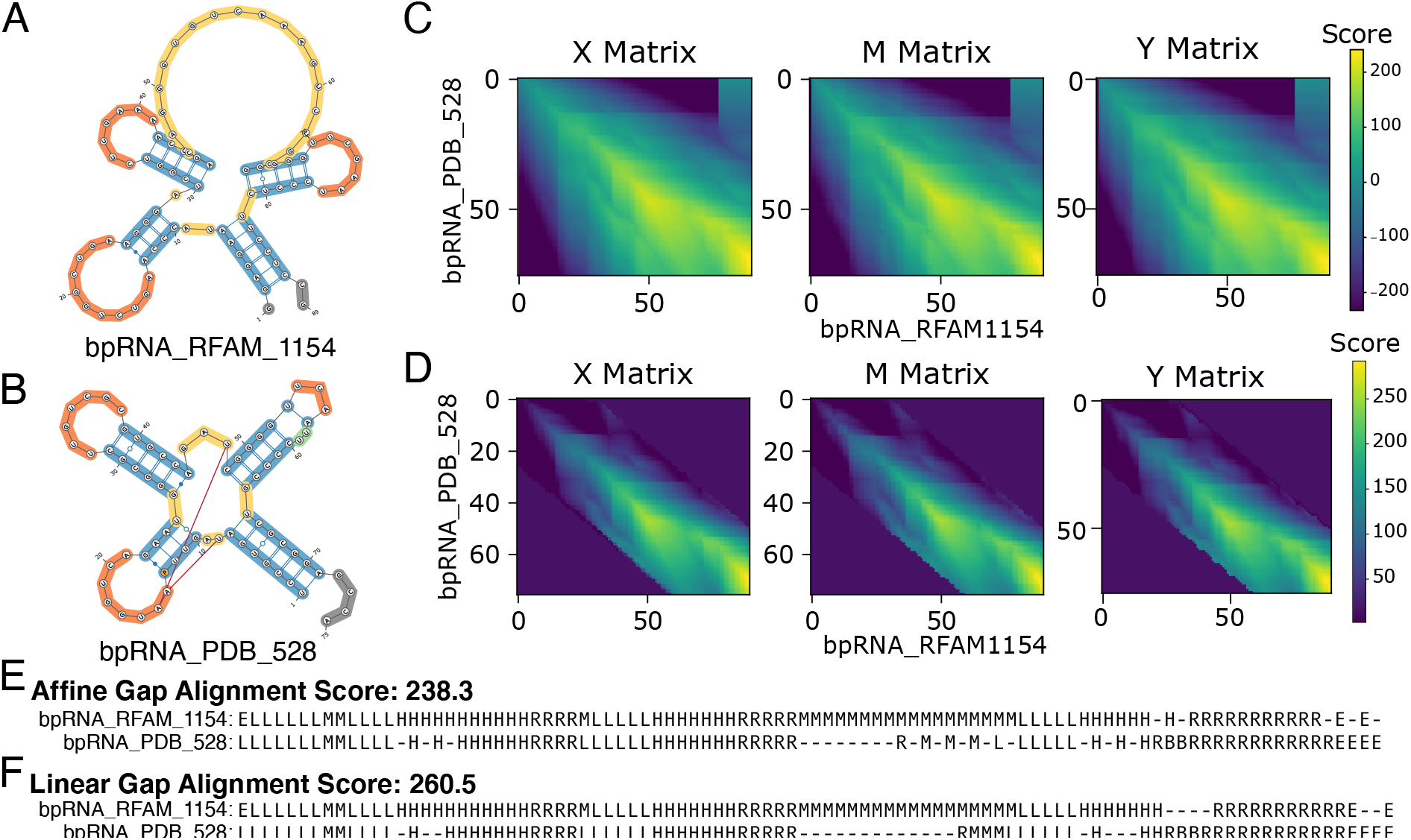
Alignment example between two tRNA structures. **A-B**. Structures bpRNA_PDB_528 and bpRNA_RFAM_1154 pulled from the bpRNA-1m database and aligned with bpRNA-align. **C**. Alignment matrices for the alignment between the structures shown in A and B, without the banded alignment customization. **D**. Alignment matrices for the alignment between A and B, demonstrating the banded alignment customization. For **C-D** negative values were mapped to zero for visualization purposes. **E**. Alignment results for the alignment of A and B without the baned alignment customization. **F**. Alignment result for the aligment of A and B with the banded alignment customization.

**Fig 3:**
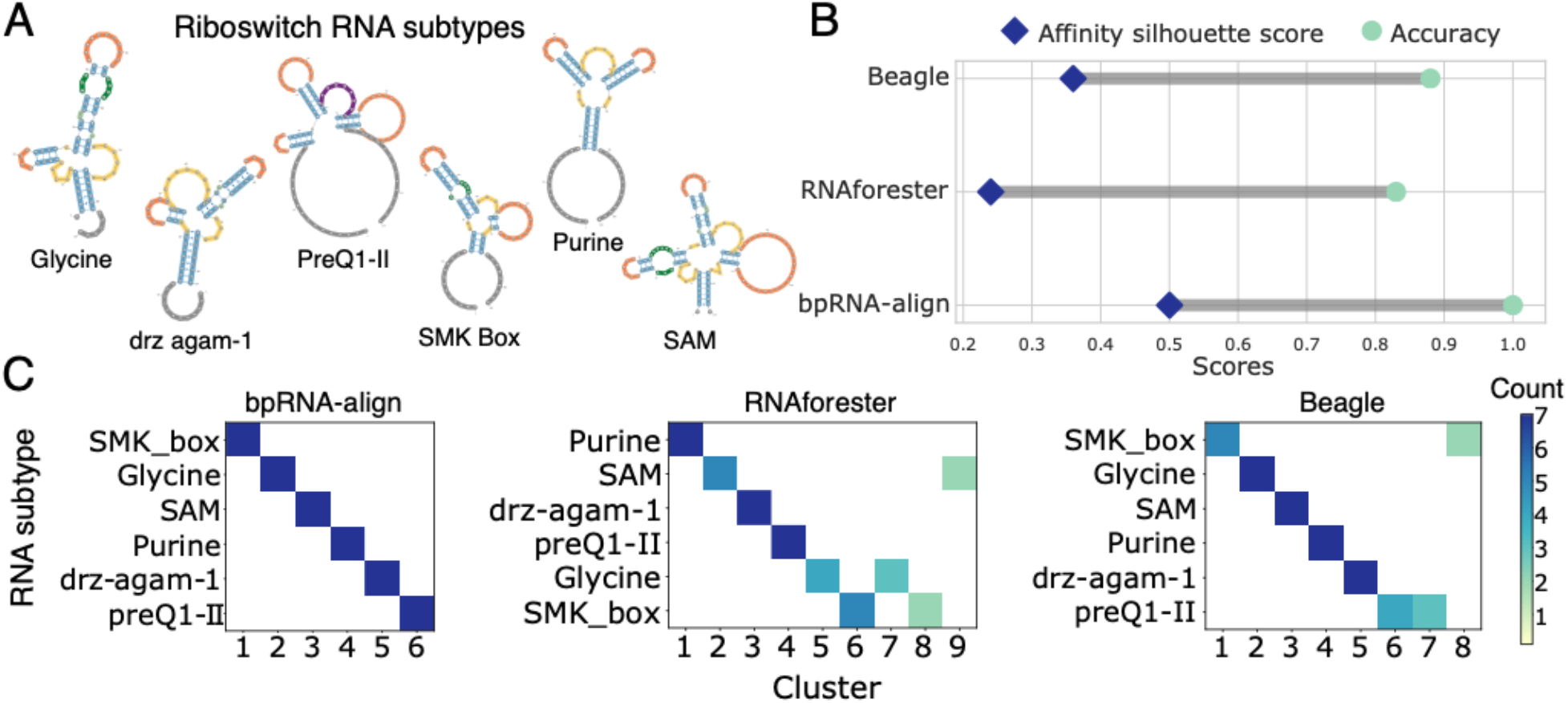
Comparison of structure comparison approaches applied to riboswitch structures. **A**. Riboswitch subtypes utilized in development of the dataset. **B**. Silhouette scores for the fixed and variable cluster number conditions resulting from each structure comparison method. **C**. Confusion matrix results for the methods compared when using a variable number of clusters. **D**. Confusion matrix results for the compared methods when using a fixed number of clusters.

### Method Comparison

To evaluate the performance of bpRNA-align, we compared it with two other RNA-comparison methods available, BEAGLE [9] and RNAforester [6]. Similarity scores were generated for each dataset using each of the three comparison methods. Clustering was performed using scikit-learn affinity propagation [17–19]. This allowed us to compute the clustering directly using the affinity scores determined from each method tested. Through clustering we collected quantitative affinity silhouette scores using the SNFpy package [20], and confusion matrices [21,22] for each method along with the corresponding accuracy. The silhouette score depicts the level of separation between clusters, with a high score resulting from easily distinguishable clusters and a low score for clusters not easily separable [20]. Confusion matrices provide visualization of the RNA families in comparison to the clustering results, where a perfect diagonal trend demonstrates a 100 percent accuracy. For each alignment method, scores were generated for all unique combinations of structural alignments within each dataset. The affinity silhouette scores, accuracies, and confusion matrices were computed to evaluate each method.

## Results

### Needleman Wunsch structure alignment

To demonstrate the functionality of bpRNA-align, we applied it on an example involving two tRNA structures with bpRNA-1m identifications of bpRNA_RFAM_1154 (Fig 2A), and bpRNA_PDB_528 (Fig 2B). These structures were chosen from different originating databases, with each structure being obtained through different methods. Structure bpRNA_PDB_528 (source ID: PDB 3TRA) originated from the PDB and was obtained through X-ray diffraction [23]. Structure bpRNA_RFAM_1154 (source ID: RFAM RF00005_AF008220.1_6334-6422) was obtained from the RFAM database and was determined through comparative sequence analysis [24]. The *X, Y*, and *M* matrices for this alignment (Fig 2C), representing the scores for all possible moves, show a strong high scoring diagonal trend, indicating the similarity between these two tRNA structures. The alignment results (Fig 2E), obtained through tracing back the ideal path between the 3 matrices, demonstrates a high similarity between the two structures with most gaps originating in the 3^rd^ branch of the multiloop.

### Banded alignment

When we applied the banded strategy to align the same two tRNA structures, with a bandwidth of *w* = 25, we obtained the same alignment result (Fig 2E). Although the same result was reached, the complexity of our method decreased from 𝒪(*n*^2^) to 𝒪(*wn*). The search space band can be visualized in the scoring matrices, *X, Y*, and *M* (Fig 2D).

### “Inverted” Affine gaps

To allow for some flexibility between structures, the affine gap strategy was implemented. Opposite of traditional sequence alignment approaches, a small gap opening cost and a greater gap extension cost was applied, which resulted in higher scores for structures with many small gap-regions throughout the structure, and lower scores for structures with long continuous gap-regions. To examine this application on an example case, bpRNA-align was implemented with both the linear gap penalty and the affine gap penalty for the pair of tRNAs in Figure 2. The main distinction between the two structures is a 15-length difference in the 3^rd^ branch of the multiloop, also known as the variable loop (Fig 2A-B). The alignment utilizing affine gaps resulted in 24 gaps and a similarity score of 238.3 (Fig 2E). In comparison, with the application of linear gaps that resulted in 26 gaps, but with a similarity score of 260.5 (Fig 2F). Therefore, the affine gap was able to produce a structural alignment using fewer gap characters, but more gapped regions. The lower similarity score when applying affine gaps, penalizes the large difference in the length of the variable loop. While we do recognize that variable loops in tRNAs can vary in length, this example illustrates how bpRNA-align penalizes large changes in loop length as opposed to small changes.

### Natural variability within structural feature types

Evaluation of the MAD of the length of each instance of a structural feature across RNA families (Fig 5A) revealed the feature types that demonstrate greatest length-MAD. Hairpins by far show the greatest length-MAD overall, with some values falling high above the mean. End and external loops also show a higher level of length-MAD, as these do not significantly affect the core structure. Stems resulted in the lowest overall length-MAD spread, with bulges, internal loops and multiloops falling close behind. Mean MAD values for each feature over all RNA families are as follows: 1.86 (Hairpin), 0.58 (End), 0.13 (Bulge), 0.14 (Stem), 0.16 (Internal loop), 1.16 (External Loop), 0.23 (Multiloop).

### Context-specific gap penalties

The addition of context-specific affine gap penalties helped improve the performance of bpRNA-align by increasing the accuracy in most of the benchmark data sets (Fig 5B). To separate out the effect of affine gaps from context-specific gaps, the accuracies of bpRNA-align with 3 different implementations was determined (Fig 5B). First, bpRNA-align was implemented with linear gaps (Linear), next it was implemented with affine gaps (Affine), and last it was implemented with context-specific affine gaps (Contextual Affine). Figure 5B shows the accuracy results for each of these cases. The addition of affine gaps helped improve the accuracy in the Riboswitch dataset only, whereas the addition of context-specific gaps improved the accuracy in 3 of the 4 datasets.

### Global alignment

Not only is our method capable of globally aligning shorter RNA structures (Fig 2) but it is also capable of accurately aligning long RNA structures. We applied and evaluated our method on both 16S and 23S structures (Supplementary Figure 1-2), ranging in length up to 3000 nucleotides. For these structures we were able to identify the regions of similarity and regions which varied from structure to structure.

### Method Comparison

The performance of the methods bpRNA-align, RNAforester, and BEAGLE were all evaluated on the 4 benchmark datasets generated from the meta-database bpRNA-1m [11]. The first dataset consisted of RNA riboswitches, the second consisted of microRNAs (miRNAs), the third consisted of three-segment structures, and the fourth consisted of four-segment structures.

### Clustering performance on riboswitch RNA

Evaluation of the Riboswitch dataset resulted in bpRNA performing the highest, with an accuracy score of 1.0 and a silhouette score of 0.50 (Fig 3B). RNAforester and BEAGLE demonstrated lower accuracies of 0.83 and 0.88 respectively, and lower silhouette scores of 0.24 and 0.36 respectively (Fig 3B). Confusion matrices provide a visualization of the number of clusters predicted and which RNA families are not correctly clustered (Fig 3C). bpRNA-align was the only method that correctly predicted the number of clusters within the riboswitch dataset. BEAGLE predicted an extra two clusters by splitting RNA subtypes SMK_box and preQ 1-II into two independent clusters. RNAforester performed the lowest, by predicting 3 extra clusters and splitting RNA categories Glycine, SMK_box and SAM each into two separate clusters.

### Clustering performance on microRNAs

Evaluation of the microRNA dataset revealed that both bpRNA-align and BEAGLE scored perfect accuracy of 1.0 (Fig 4B), with BEAGLE obtaining a higher silhouette score (0.59) compared to bpRNA (0.33) (Fig 4B). Assessment of the confusion matrices showed that RNAforester did not correctly cluster the miRNA subtypes, resulting in a lower accuracy score (0.89) and lower silhouette score (0.11) (Fig 4B). Clustering of RNAforester resulted in miss-grouping of the RNA subtypes mir-166, mir-286 and mir-159 by either splitting into multiple independent clusters or combining incorrect categories together in a cluster (Fig 4C).

**Fig 4:**
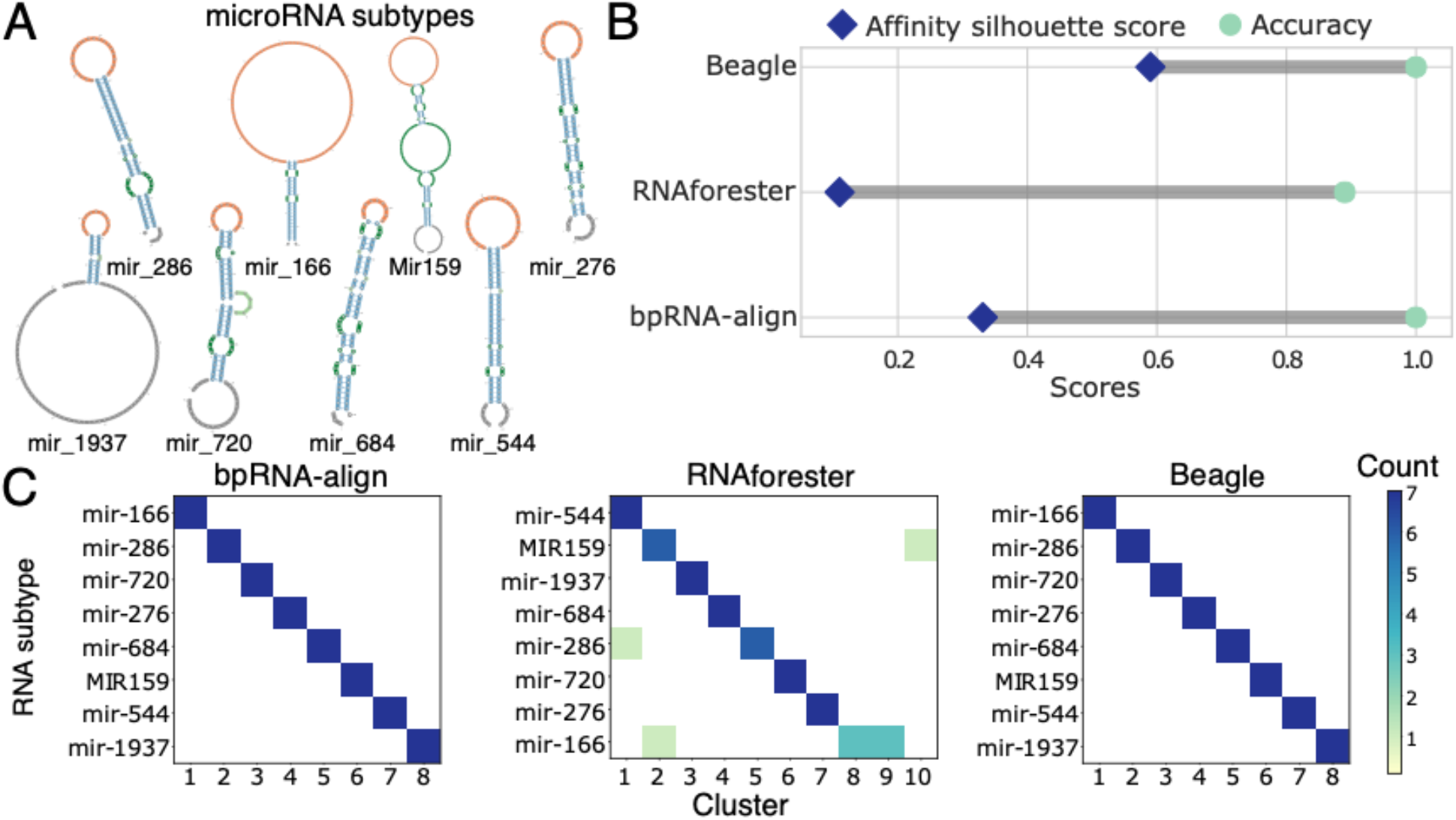
Comparison of structure comparison approaches applied to micro-RNA structures. **A**. Micro-RNA subtypes utilized in development of the dataset. **B**. Silhouette and accuracy scores resulting from each structure comparison method. **C**. Confusion matrix results for the methods compared.

### Clustering performance on three-segment structures

The three-segment structure dataset was the only example where bpRNA-align did not achieve or match the highest accuracy. Clustering resulted in RNAforester obtaining 1.0 accuracy, followed by bpRNA-align, 0.95, and BEAGLE, 0.89 (Supplementary Figure 3B). Silhouette scores for RNAforester, bpRNA-align and BEAGLE are determined to be 0.33, 0.42 and 0.51 respectively (Supplementary Figure 3B). Evaluation of the confusion matrices (Supplementary Figure 3C) show that both BEAGLE and bpRNA-align split RNA family ROSE into two independent clusters. BEAGLE also splits RNA family His_leader into two clusters.

### Clustering performance on four-segment structures

For the dataset consisting of four-segment structures, RNAforester and bpRNA-align predicted accuracies of 1.0, with BEAGLE obtaining accuracy of 0.93 (Supplementary Figure 4B). Silhouette scores for the three methods were 0.42 for RNA forester, 0.54 for bpRNA-align and 0.58 for BEAGLE (Supplementary Figure 4B). Visualization of the confusion matrices (Supplementary Figure 4C) shows that BEAGLE was not able to correctly predict the RNA family BTE, where it clustered the structures into two independent clusters.

## Discussion

### How bpRNA-align addresses feature-specific length variability

To compare and cluster RNA structures, there needs to be a metric capable of accurately representing the similarity between two structures. This metric should consider the natural variation displayed within each structural feature over a broad range of RNA families. This requires that the structure representation used to compare structures can distinguish different types of unpaired regions. Our method utilizes a compact yet descriptive structure representation which, unlike RNAforester, differentiates between unpaired regions RNA structure, and distinguishes between left- and right-handed stems using a compact alphabet, unlike BEAGLE.

To address the variability in feature length, bpRNA-align utilizes an inverted, context-specific affine gap strategy, which not only allows for differentiation between gap openings and gap extensions, but also allows for gap penalties to depend on the feature type. Context-specific gap penalties have been used previously in protein alignment to better capture the natural variation in constraints on protein structure [25]. Our use of context-specific gap penalties improved performance, and was guided by the natural variation in the lengths of RNA structural features. These gap penalties are calculated and weighted by the observed mean MAD for each structural feature. By applying lower gap penalties in feature types with more variation, and higher gap penalties in feature types with less variation, our method provides the adaptability to correctly capture structural similarity. Our analysis of the MAD length for each structural feature over 26 different RNA categories revealed a broad range of variation (Fig 5A). Our analysis shows that the feature types of stems, multiloops, bulges and internal loops should be weighted heavier in the scoring process and is a consideration we took when developing bpRNA-align.

**Fig 5:**
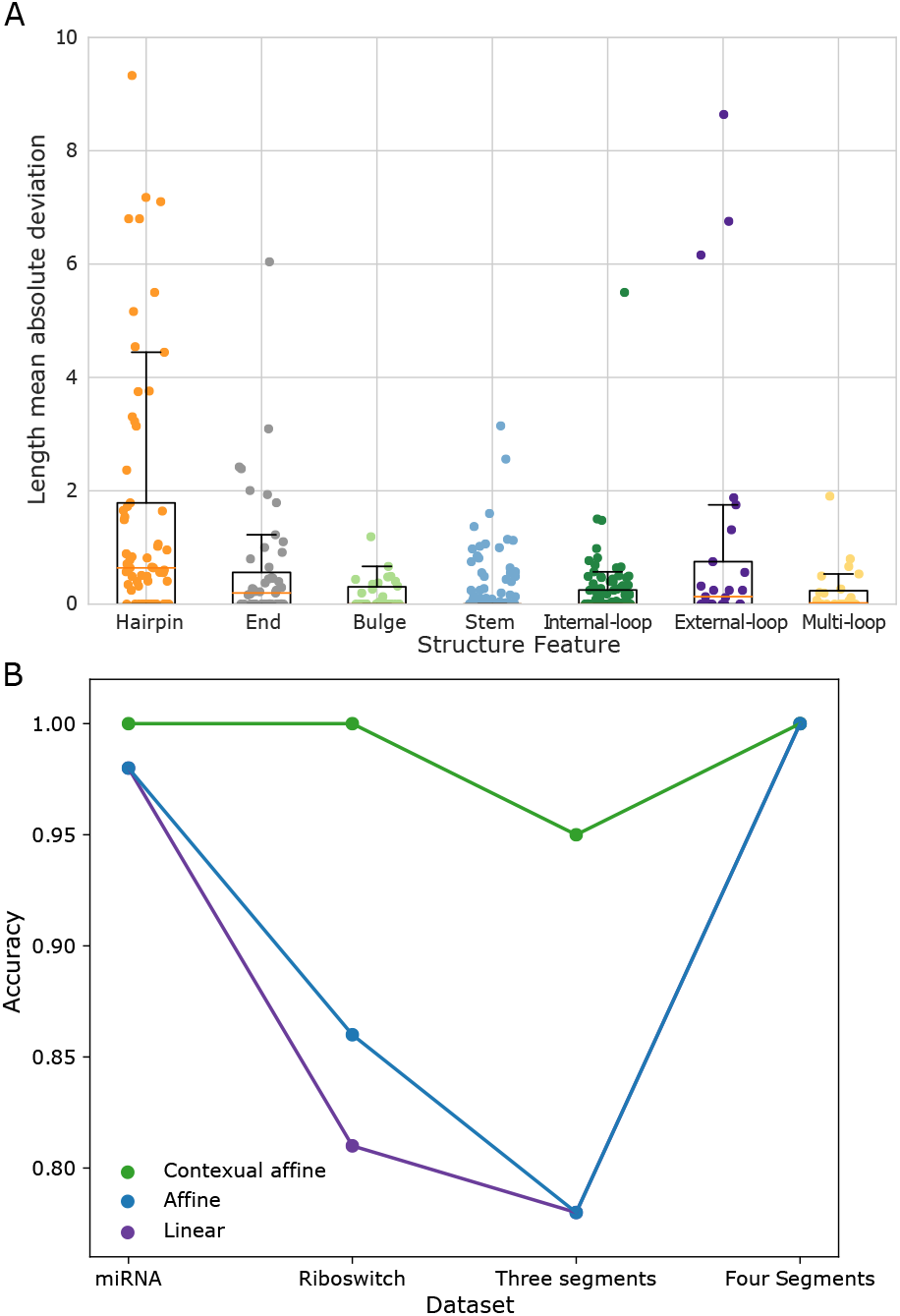
Utilization of the mean absolute deviation of length to guide gap penalties. **A**. Feature specific box plot of MAD values for each character in the feature pattern of each RNA subtype. **B**. Comparison of bpRNA-align before the addition of context-specific affine gaps (purple), bpRNA-align with only affine gaps implemented (blue), optimized bpRNA-align (dark blue) with context-specific affine gaps.

### Benchmarking bpRNA-align through biological clustering

We wanted to evaluate our methods ability to identify RNA functional categories. An ideal approach for this is a type of biological clustering [26]. This type of framework allows for evaluation of the alignment quality based on its ability to reach the biological goal (grouping RNAs from the same category) and not just a comparison on the sequence level [27]. To achieve this, we sought to develop benchmark datasets composed of different RNA families to demonstrate the accuracy of each methods ability to capture the biological context. The four datasets developed in this work represent similar structures from different RNA categories, creating challenging cases to cluster, based on similarity scores. To determine the accuracy, affinity propagation was applied to cluster the datasets. This is an ideal method, as it is based on the similarity scores from each method to perform clustering rather than a distance.

### Assessing method performance and scoring

bpRNA-align matches or outperforms the other methods in 3 out of the 4 datasets examined in this work. This is likely due to the type of structural representation used and the inherent adaptability added within this method, which allows it to be more robust over a broader range of RNA structures. The inverted affine gap implementation with a small gap opening allows for flexibility in the alignments without leading to large separations between similar structures. A large portion of this flexibility is also due to the addition of context-specific gap opening and extension penalties. This allows for flexibility in structure features that are less stable and more likely to vary in length.

RNAforester demonstrates large scoring effects for gaps, but mismatches in structure features appear to have little consequence on the score. An example low penalty for mismatches resulting in different RNA types scoring highly is the alignment between bpRNA_RFAM_6133 of category mir-166 and bpRNA_RFAM_18555 of category MIR159 (Supplementary Figure 5). Although these structures appear completely different patterns of base-pairing, they are scored much higher than alignments of the same RNA type in the mir-166 category. In other cases, RNAforester split a single RNA family into multiple clusters due to large section of gaps in variable-length regions, which leads to a large differentiation between structures of the same RNA type. For example, hairpins are observed to have a large level of variation (e.g. mir-166, Supplementary Figure 5) in many RNA families, and in these cases RNAforester scores may be too sensitive to cluster the families correctly.

Within the datasets analyzed in this work, BEAGLE is observed to predict too many clusters in 3 of the 4 datasets examined. This result is due to splitting RNA families into multiple clusters. To understand why BEAGLE was splitting categories into multiple clusters, each split family was examined individually. This revealed that the number of gap characters within each alignment varied largely between a single RNA category, most often within an unpaired region that has more variability. As a result, BEAGLE has a larger scoring distinction between certain structures within the same family, and this results in splitting the family into multiple clusters.

### Future directions and considerations

An aspect that both RNAforester and BEAGLE apply in their alignment approach is the use of primary sequence and not just structure alone. This is something bpRNA-align does not currently account for but is an addition we plan to make in the future. We know that some types of RNAs are more conserved in their primary sequences than others, and thus it is important to apply a user-weighted adjustable option for sequence information. This could potentially boost the performance of the alignment results, especially for cases that are less structurally conserved, and more sequence conserved.

Another factor that could boost performance is an optimized substitution matrix. bpRNA-align currently utilizes a knowledge based phenomenological substitution matrix, but there is room for improvement here. The alignment substitution matrix can be improved using the log-likelihood method, analogous to the generation of the BLOSUM [28] and point accepted mutation (PAM) substitution matrices for amino acids [29] but instead with RNA structural elements. BEAGLE constructs a log-likelihood substitution matrix, but as mentioned previously its use of different characters for each loop length results in a large number of parameters to calculate. Our approach will rely on a set of high-similarity structural alignments, curated from known homologous RNAs. By examining the empirical frequency of each type of substitution, the structural substitution matrix can be computed.

