## Supplementary Figures for "bpRNA-align: Improved RNA Secondary Structure Global Alignment for Comparing and Clustering RNA Structures"

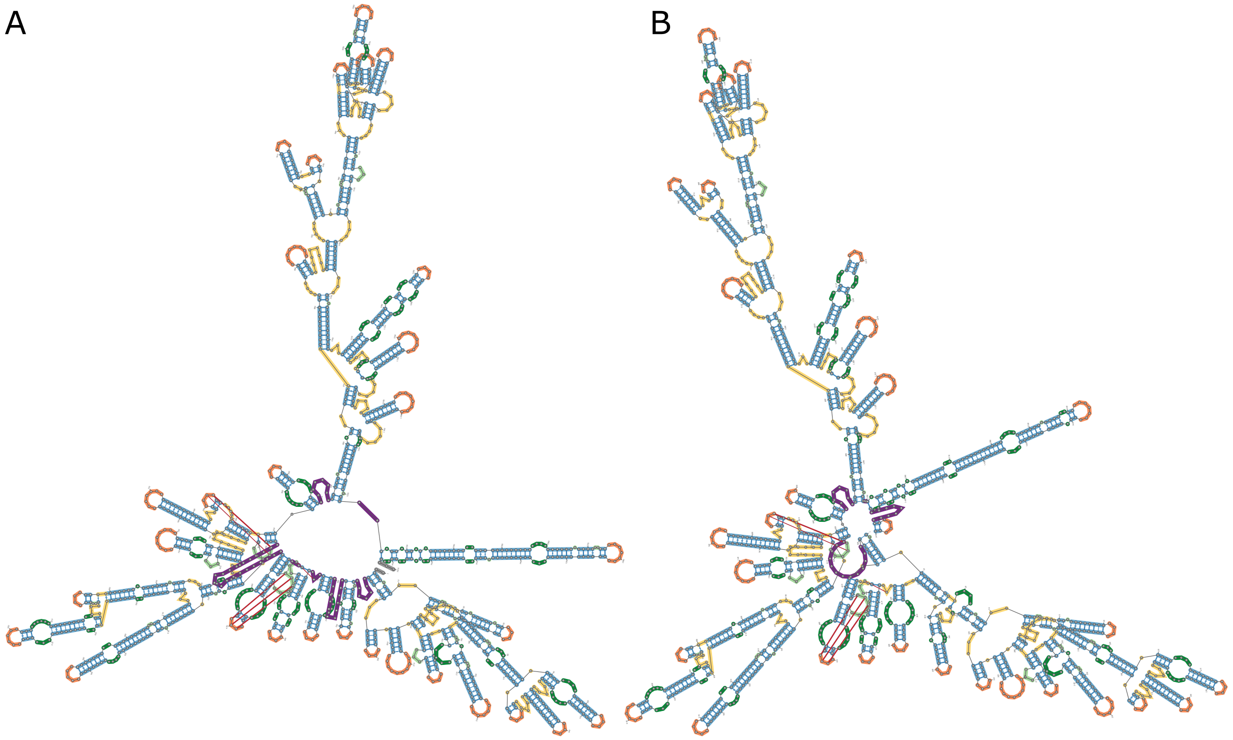


Structure 1: bpRNA_CRW_2 Structure 2: bpRNA_CRW_29

C

```
1: EELLLBLLLM-M-L-L-L-B-B-L-L-L-L---L-B-L-L-L-LL-B-LL-HHHH-RRRRRRRRBRRRRRRMMMMMLLLMMMMMLLLLLLLBBLLIIILLLLLLLMLLLIIIILLLLLLLHHHHRRRRRRRIIIIIRRRRMMMMLLLLLLLLLHHHHRRRRRRRRRMMMMLLHHHHRRM
2: LBLLL-LLLMLLLLLLLLLIIIILLLBLLLMMLLLBBLLLLLBLLLLLLLLHHHHRRRRRRRRRRBRRRRRRMMMMMLLLMMMMMLLLLLLLBBLLIIILLLLLLLMLLLIIIILLLLLLLHHHHRRRRRRRIIIIIRRRRMMMMLLLLLLLLLHHHHRRRRRRRRRRMMMMLLHHHHRRM
1: RRRRRRRIIRRRRRRRRRRRLLLBBLILLLBBLLLLLLLLHHHHHHHRRRRRRRRRRRIIIIIRRRRMMLLLLLLLLHHHHRRRRRBBBRRRRRRRMLLLLLLHHHHHHHHHHHRRRRRRRMLLLLHHHHRRRRRMMMRRRRRRXXXXXXXLBLLLIIILLLBLHHHH-RRRRIRRRRXXXXXXXXXXXXXLLL
2: RRRRRRRIIRRRRRRRRRRRLLLBBLILLLBBLLLLLLLLHHHHHHHRRRRRRRRRRRIIIIIRRRRMMLLLLLLLLHHHHRRRRRBBBRRRRRRRMLLLLLLHHHHHHHHHHHRRRRRRRMLLLLHHHHRRRRRMMMRRRRRRIRMMMMMLBLLLIIILLLLLHHHHRRRRRIRRRRMRRBRRRRRRMMLLL
1: LIIIIIILLLLHHHHRRRRIIIIIRRRRXXXLLLLLLIIILIILLHHHHRRIIRIIIIRRRRRRBBBRXXLLLLLBBBBBBLLLLLIIIIILLHHHHRRIIIIIIIIRRRRRRRRRRXXXXXXXXXXXXXXXXXXXXXLLLBBBBBLMLLLLILBLLLMLLLLLLBLLLLLLLLLLLIIIILLLLLLL
2: LIIIIIILLLLHHHHRRRRIIIIIRRRRMMMLLLLLLIIILIILLHHHHRRIIRIIIIRRRRRRBBBRMMLLLLLBBBBBBLLLLLIIIIILLHHHHRRIIIIIIIIRRRRRRRRRRMRRRRRRXXXXXXXXXXXXXLLLBBBBBLMLLLLILBLLLMLLLLLLBLLLLLLLLLLLIIIILLLLLLL
1: LHHHHHRRRRRRRIIRRRRRRRRRRBBRRRRRRRRRRMMMLLLLLLLLLIILLLLLLLLLIILLLLLLLLLIIILLLHHHHHHRRRIIIIIIRRRRRRRRIIRRMMMMLBBLLHHHHRRRRRRRRRRRIRRRRRRBRRRMMRRRRIIRRRRMMMMLLLLLLBBBLLIIILLLLHHHHHHHHHRRRRRIIIRR
2: LHHHHHRRRRRRRIIRRRRRRRRRRBBRRRRRRRRRRMMMLLLLLLLLLIILLLLLLLLLIILLLLLLLLLIIILLLHHHHHHRRRIIIIIIRRRRRRRRIIRRMMMMLBBLLHHHHRRRRRRRRRRRIRRRRRRBRRRMMRRRRIIRRRRMMMMLLLLLLBBBLLIIILLLLHHHHHHHHHRRRRRIIIRR
1: RRRRRRMMMMMMMMMMMLLLLLMMLLLLLLLLLLLLHHHHHHHRRRRRRRRRRRRMMMLLHHHHRRMMMMMRRRRRRRRRRXLLLIIIIIILLLLHHHHRRRRIIIIRRRXXXXXXXXLLLLLBLLLLLLLILLMMLLLLLLLLLLLLLLLLLLMMMMLLLHHHHHHHHHRRRRMMMMMMMMMLLLLLLL
2: RRRRRRMMMMMMMMMMMLLLLLMMLLLLLLLLLLLLHHHHHHHRRRRRRRRRRRRMMMLLHHHHRRMMMMMRRRRRRRRRRXLLLIIIIIILLLLHHHHRRRRIIIIRRRXXXXXXXXLLLLLBLLLLLLLILLMMLLLLLLLLLLLLLLLLLLMMMMLLLHHHHHHHHHRRRRMMMMMMMMMLLLLLLL
1: MMMMMLLLLLLBBLMMLLLLLLLHHHHRRRRRRR-MMLLHHHHRR-M-RRRRRRR-MLLBLLLLBBLLLLLLLLLMMMLLLLLLLLLHHHHRRRMMLLLLHHHHHHRRRRMMRRRRRRMMMMMLLLLMLLLLLLLIIIILLBBLLLHHHHHHRRRRRIIIIRRRRRRRMMLLLLLLLHHHHHHRRRRRR
2: MMMMMLLLLLLLLLMMLLLLLLLHHHHRRRRRRRMMMLLHHHHRRMMRRRRRRRRRMLLBLLLLBBLLLLLLLLLMMMLLLLLLLLLHHHHRRRMMLLLLHHHHHHRRRRMMRRRRRRMMMMMLLLLMLLLLLLLIIIILLBBLLLHHHHHHRRRRRIIIIRRRRRRRMMLLLLLLLHHHHHHRRRRRR
1: RRMMMMMRRRRMMMMMRRRRBRRRBBBRRRRRRRRRMMMMRRRRRRRMMMRRBRRRRRRRRRRMMMLLLLLLIIILLLLLIILLIIILLLHHHHRRRIIIRRIIIIRRRRRIIIRRRRRRRMMMMMLIIILLLLLLLLLHHHHHHHHHRRRRRRRRIIIRMMMMMRRRRRMMMMMLLLLLLLLHHHHHHH
2: RRMMMMMRRRRMMMMMRRRRBRRRBBBRRRRRRRRRMMMMRRRRRRRMMMRRBRRRRRRRRRRMMMLLLLLLIIILLLLLIILLIIILLLHHHHRRRIIIRRIIIIRRRRRIIIRRRRRRRMMMMMLIIILLLLLLLLLHHHHHHHHHRRRRRRRRIIIRMMMMMRRRRRMMMMMLLLLLLLLHHHHHHH
1: HRRRRRRRRMMMMMMRRIIIRRRRRRRRRRRBRRXXLILLILLILILLLLLLLLLIILLLLLLLLLLLLLIIIILLLLLLBLLLLBBLLHHHHHHHRR-RRRRRRRRRRIIIRRRRRRRRRRBRRRIIRRRRRRRRIIRIRRIIRRIIR----E-E-E--------
2: HRRRRRRRRMMMMMMRRIIIRRRRRRRRRRRBRRXXLILLILLILILLLLLLLLLIILLLLLLLLLLLLLIIIILLLLLLBLLLLIILLHHHHHHHHRRIRRRRRRRRRRRRIIIRRRRRRRRRR-RRRIIRRRRRRRRRIIRIRRIIRRIIRXXXXXXXXXXLLHHHHHRR
```

Supplementary Figure 1: **bpRNA-align results for aligning two 16S structures from the bpRNA-1m meta database. A.** 16S RNA structure with ID bpRNA_CRW_2. **B.** 16S RNA structure with ID bpRNA_CRW_29. **C.** Alignment results between the two 16S structures.

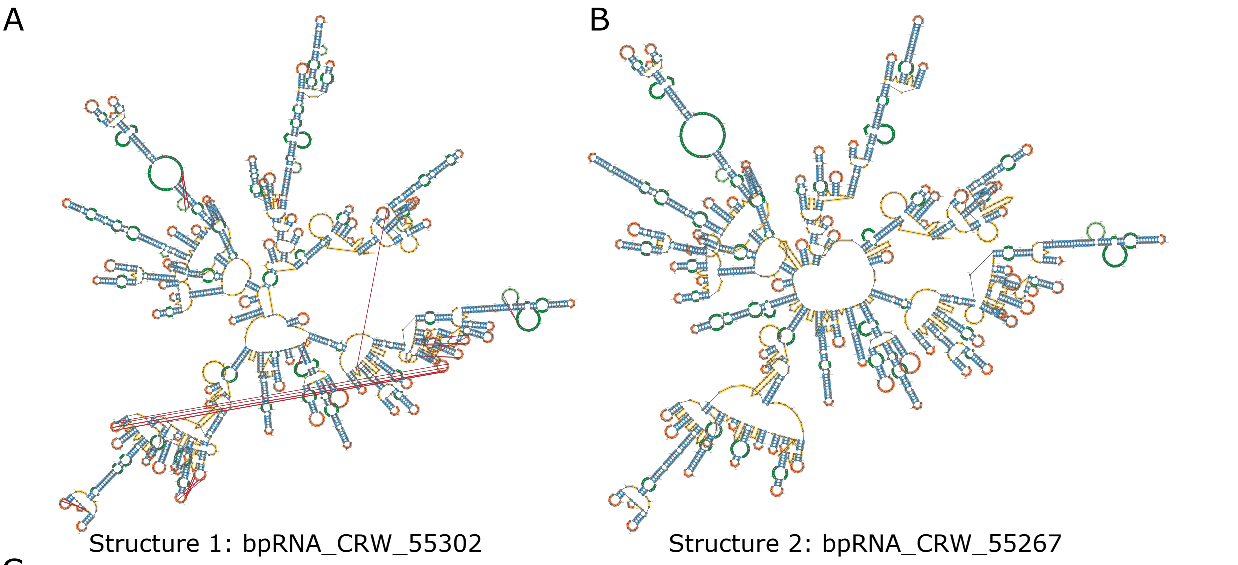


C

```
1: -LLLLLLLLMMMMMMLLLLLLLLLLIIIIILLLMMLLLLLLLLLLLLLLMMMMMLLLLLLLHHHHHHHHHRRRMMMMMLLLLLLLIILLLHHHHHHHHRRRIIIIIRRRRRRRMMMRRRRMMMLLLHHHHRRRLL
2: LLLLLLLLLMMMMMMLLLLLLLLLLIIIIILLLMMLLLLLLLLLLLLMMMMMMLLLLLLHHHHHHHHHRRRMMMMMLLLLLLIIILLLHHHHHHHHRRRIIIIIIRRRRRRRMMMRRRRMMMLLLHHHHRRR-L
1: LLLLLLHHHHRRRRRRRRRMLLLLLLLLLHHHHHHHHHHRRRRRRRRRMRRMMMMMLLLLLLIIIIILLHHHHRRIIIIIIRRRRRRBRMMMMMMMMMMLHHHHRRMMLLLLLLIIIIIILLHHHHHHRRIIIIRRRR
2: -LLLL-HHHH-R-RRRR-M-L-L-LLL-H---H-H-H-R-R-RRR-M-MMMMMLLLLLLIIIIILLHHHHRRIIIIIIRRRRRRBRMMMMMMMMMMLHHHHRRMMMLLLLLIIIIIILLHHHHHHRRIIIIRRRR
1: RMMLLLLMMMM--L--L-L-L-LL-HHHH--R-R-R-R----RR-MMMMMM-LLLLLLLLLIIIIILLIIILLLL-LLLLLL-I-LLLMMML-LLLLHHHHHHHRRRRRMMLHHHRMLLLHHHHHHHHHRRRMMM
2: RMMLLLLMMMMLLLIILLLLLLLLLLHHHHRRRRRRRRRRIIIRRRMMMMMMMLLLLLLLLL-L-BBLL-L-LLLLBLLLLLLLIIILLLMMMLBLLLLHHHHHHHRRRRRMMLHHHRMLLLHHHHHHHHHRRRMMM
1: RRRIII-RRRRRRRRRRIIRRIIRRRRRRBBRRRRMMMMMLLLLLLLL-HHHHHHHHRRRRRRRRRMMMMMMLLLLL-HHHHHHRRRRRMMRRRRRMMRMMRRBRRRRRRRBRRMMMMMMMMMMMMMMMMLLHHHHR
2: RRRIIIIRRRRRRRRRRR-RRR-RRRRRRRB-RRRMMMMMMMLLLLLLHHHHHHHH-RRRRRRRMMMMMMLLLLLHHHHHHHHRRRRRMMRRRRRMMMMMRR-RRRRRRRBRRMMMMMMMMMMMMMMMMLLHHHHR
1: RMMMMRRMMMMMMMMMMLLLL-HHHHRRRRMMMMMMMMMMMMMMRIIIIIRRRRRRRRRR-M-MMMM-L-M-L-L-L-----L------L-L-L-L---L-----L-L-H-HH--H-R-----R---R-R-R–
2: RMMMMRRMMMMMMMMMMLLLLHHHHHHRRRR-MMMMMMMMMMM-RIIIIIRRRRRRRRRRRRMMMMMMMLLLLLLLLLLIIIILLIIIIIIILLLLLLLIIILIIIILLLLLHHHHRRRRRIIIIIRIIRRRRRRRI
1: ---R-R-R-R-R-R-MMLLLLHHHHHHHHHHRRRRLLLLLLMMMMLLLLLLLLLLLLLLMMMLLLLIILLL-HHHHHHRRR-IIRRRRRMMMLLHHHHHRR---M-L-LL---L-L--H-H----------------
2: IIRRIRRRRRRRRRRRRMMLLLLHHHHHHHHHHRRRRLLLLLLMMMMLLLLLLLLLLLLLL-MMLLLLIILLLLLHHHHHHRRRRIIRRRRRMMMLLHHHHHRRMMMMMLLLLIIILLLLHHHHHRRRRIIRRRRRMMLLLL
1: -H-HHRRRRRRMMM-RRRRRRRRRRRRRRMMLLILIIILLLLLLMMMLLLLLLLLLLLLLMLLLLIIILLLLLLLHHHHHRRRRRRRIIIIRRRRMMM-LLLBL-LL-LLHHHHHHHHHRRRRRRRRRMMRBRRRRRRR
2: LHHHHRRRRRRMMMMRRRRRRRRRRRRRRMMLLILIIILLLLLLMMMLLLLLLLLLLLLLMLLLLIIILLLLLLLHHHHHRRRRRRRIIIIRRRRMMMMLLL-LBLLBLLHHHHHHHHHRRRRRRRRRMMRBRRRRRRR
1: RRRRRMLLLHHHHHHRRRMMMMMMRRRRRRIIIIIIRIRRMMLLLLLLMMMMLLLLHHHHHRRRRRMLLLLLLLBBBLLL-LLLLLLLLLIIIILLLIII-I-LLLILLLLIIILLLLLLLHHHHHHRRRRRBR
2: RRRRRMLLLHHHHHHRRRMMMMMMRRRRRRIIIIIIRIRRMMLLLLLLMMMMLLLLHHHHHRRRRRM-LLLLLLLIIILLLBLLLLLLLLIIIILLLIIILLLLLLLLLLLLBB-LLLLLLH-HHHRRRRRRR
1: RIIIRRRRIRRRIIRRBRIIIIRRRRRRRRRR-B-R-R-RBRRRRRBRMMMLLLLLLLLLLHHHHHHRRRRRRRRRRRMMMMLLHHHHHHHRRRMMMLLLBLLLIIILLLBBBBBBL-LLLLLL-I-I-I-I
2: R-RRRRRRRRRRIIRRBRIIIIRRRRRRRRRRRIIIIIIR-RRRRRBR-MM-LLLLLLLLLHHHHHH-RRRRRRRRRRMMMMLLHHHHHHHRRRMMMLLLL-LLLIIILLLBBBBBBLBLLLLLLIIIIIIII
1: IIIIIII-LLLBLLLLLLLLLLIIIIIIILLLLLMLLLIILLLHHHHHHHHHHHRRRIIRRRLHHHRLIIILLHHHHHHHRRIRMRRRRRIIIRRRRRRRRRRRRRRIIIIIIIIIIIIIIIIIIIRRRRRRRRRR
2: IIIIIIIILLLBLLLLLLLLLLIIIIIIILLLLLMLLLIILLLHHHHHHHHHHHRRRIIRRRLHHHRLIIILLHHHHHHHRRIRMRRRRRIIIRRRRRRRRRRRRRRIIIIIIIIIIIIIIIIIIIRRRRRRRRRR
1: IIIRRRRRRR-L-LLLLLL-HHHH-R-RRRRR--R-MMMMRRRRRRLILLLLLLLLLLIIIIIILLLLLLLHHHHRRRRRRRRRIIIRRRRRRRRRRRIIRMMMMMRRRRRR-L-L-IIIIILLMMMLLLLLL
2: IIIRRRRRRRLLLLLLLLLLHHHHRRRRRRRRRBBRRMMMMRRRRRRLILLLLLLLLLLIIIIIILLLLLLLHHHHRRRRRRRRRIIIRRRRRRRRRRRIIRMMMMMRRRRRRMMMMMMMMMMMMMMMLLLLLL
1: HHHHHHHHRRRRRRLLLLMMMMLLLLMMLLMMMLLLLLLIIILLHHHHHHRRIRRRRRRRMMMMMMLLLLMLLIIILLLLILILLL-HHHHRRRI-RIRRRIII-RRMMMLLLLHHHHHHHHHRRRRRMM-L-LLL
2: HHHHHHHHRRRRRRLLLLMMMMLLLLMMLLMMMLLLLLLIIILLHHHHHHRRIRRRRRRRMMMMMMLLLL-LLIIILLLLLLLLLLLLHHHHHHRRRBRRRRRRIIIRRMMMLLLLHHHHHHHHHRRRRRMMLLLLLL
1: LLLLL-LLLL---I----I-L-L-L-LLL-IIILLLLIILLLIILLLLLLLLLLL-HHHHHHHHHRRRRRR-LLIILLLLIII-LLLLLLLLLLLBLLHHHHHHRRRRRRRRRRRBBBBRRRRIII-RRRRIIIRR
2: LLLLLMLLLLHHHHRRRRMMLLLLLBLLLLIIILLLLLLLLLLIILLLLLLLLLLLLLH-H-H-HHRRRRRRRR-MMMLLLLIIIILLLLLLLLLLLLLLLHHHHHHRRRRRRRRRR--R-RRRRIIIIRRRR-MMMM
1: MLLLLIIILLLHHHHRRRIIIIRRRRMMRRRRBRIIIRRRIIIRRRRIIIIIIII-RRRRRRIRRBBBBBRRR-RRRRRRRRRRRRRMMRRMMMMMLLHHHHHHHRRMRRRRRMLLLL-H-HHHHRRRRRMMRRRR
2: MLLLL---LLLHHHHRRR----RRRR-MRRRR-RIIIRRR-RRRRRRIIIIIIIIIRRRRRRRRR-M-M-RRRBRRRRRRRRRRRRRMMRRMMMMMLLHHHHHHHRRMRRRRRMLLLLHHHHHHHHHRRRR-MRRRR
1: MM-LLLLIIII-LLLLLLLLILLLMMMMMMMMMMMMMMLLLLLIIILLHHHHHRRIIIIRRRRRR-L-L-LLLL-HHHHH-H-RRRRRR-LHHHRMMMMMMMLLLLLLLLMMLLLHHHHHHHHHHHRRRMMLLLL
2: MMMLLLLIIIIILLLLLLLLLLLLMMMMMMMMMMMMMMLLLLLIIILLHHHHHRRIIIIRRRRRRLLLLLLLLLLHHHHHHRRRRRRRRRRLHHHRMMMMMMMLLLLLLLLMMLLLHHHHHHHHHHHRRRMMLLLL
1: LLLLLIILLLLHHHHRRRRIIRBBRRRRRRRRRMMLLLLMLLLBBLLLLLLLLBBLLLIIIILLILLLLLLLLIILLHHHHRRIIIRRRRRRRRIRRIIIRRRRRRRRRBBRRRRRRLLLLLLHHHHHHHHRRRRRRLHH
2: LLLLLIILLLLHHHHRRRRIIRBBRRRRRRRRRMMLLLLMLLLBBLLLLLLLLBBLLLIII--I-L-L-LL--L-HHHH-H-H-R-RRRR-I--IIIRRRRRRRRRBBRRRRRRLLLLLLHHHHHHHHRRRRRRLHH
1: HRMMMMMLBBBBBBBBBLLLLLLHHHHHRRRRRRRMMMMMMMMMMRRRRMMMRRBRRRRRRRRMMMMRRRIRRRRRRRRIR-RRRRRIIIIRRRMMMMLLLLHHHHHHHHRRRRRMMLLLLLLLLBLLLILLM
2: HRMMMMMMMMMMMMMMMLLLLLLHHHHHRRRRRRMMMMMMMMMMRRRRMMMRRBRRRRRRRRMMMMRRR-RRRRRRRR-RIRRRRMMMMMMMMMMMMLLLLHHHHHHHHRRRRRMMLLLLLLLLBLLLILLM
1: MMMMMLLLLBLLLLLLLMLLLLIIILLLLLLLLMMLLLLLLLLLLLLLLLLLLLBBBBBBBBBLLLLLIILLLIIIIIILLLLLLLLHHHHRRRRRRRRRIIRRRIIIIIIIIIIIIIRRRRRRRRRRRRRRR
2: MMMMMLLLLBLLLLLLLMLLLLIIILLLLLLLLMMLLLLLLLLLLLLLLLLLLLBBBBBBBBBLLLLLIILLLIIIIIILLLLLLLLHHHHRRRRRRRRRIIRRR-IIIIIIIIIIIIRRRRRRRRRRRRRRR
1: RRRRRRRMMMLLLBLLLLLLHHHHHRRRRRRRRRMMMMMRRRRRRRIIIIRRRRMMLLLLHHHHHHRRR-RLLLLLLHHHHHHHHHHHHRRRRRRLMLLMMMMLLLLLLLLILLLLLLHHHHHHHHRRRRRRIII
2: RRRRRRRMMM-LL-L-L-L-H-H-H-R-R-RRR-MMMMMRRRRRRRIIIIRRRRMMLLLLHHHHHHRRRBRLLLLLLHHHHHHHHHHHHRRRRRRRMMLLMMMMLLLLLLLLILLLLLLHHHHHHHHRRRRRRIII
1: RRLLHHHHHHHRRMMMMRRRRRRRMMMMLLLLLLLLL-H-H-HHHHRRRRRRRRRRMLLLHHHHRRRMRRMMMMMMMMMMLLLLL-LLBLLHHHHHHHHHRRRRRRRRRRMMMMMRMMMMMMRRRRRBBRRRRRRRMM
2: RRLLHHHHHHHRRMMMMRRRRRRRMMMMLLLLLLLLLHHHHHHHHHHRRRRRRRRRRMLLLHHHHRRRMRRMMMMMMMMMMLLLLLBLLLLLH-HHHHHHRRRRRRRRR-RMMM-MMMMMMRRRRRBBRRRRRRRMM
1: MMMMMMLLLBLLLLLLLLLIIILLHHHHHHHRRIIIRRRRRRRBBRRRRMMMMMMMMMMLLLLLLLLLLLMMLLILLLLLLHHHHHHRRRRRRIIIRRMLLLLHHHHHHRRRRRMMMMMRRRRRBBRBBRRRR
2: MMMMMMLLLBLLLLLLLLLIIILLHHHHHHHRRIIIRRRRRRRBBRRRRMMMMMMMMMMLLLLLLLLLLLMMLLILLLLLLHHHHHHRRRRRRIIIRRMLLLLHHHHHHRRRRRMMMMMRRRRRBBRBBRRRR
1: BRMMMMMLLLLLLLHHHHRRRBRRRRMMMMRRIIRRRRRRRRRRRRMMMMLLLLLLLLIIILLLMMLLLLLLL-HHHHHHHHHHHHHHHHHRRRRRRR-LLLLLLIIILLLIIIILLLLLLLLLLLLHHHHRRRRR
2: BRMMMMMLLLLLLLHHHHRRRBRRRRMMMMRRIIRRRRRRRRRRRRMMMMLLLLLLLLIIILLLLMMLLLLLLLLHHHHHHHHHHHHHHHHHRRRRRRRR-LLLLLIIILLLIIII-LLLLLLLLLLHHHH-RRRR
1: RBBRRRRRIIIIRRRIRRRRRRMMLLLIIIILLLLLHHHHHHHHHHHRRRRRIIIIRRRMRRRIIIIIIIIRRRRRRRRMM-------L---L-L-L-----H-H-H-H-R-R-R-R-M-L-I-I-LLLLLLL
2: RBBRRRRRIIIIRRRI-RRRRRMMLLLIIIILLLLLHHHHHHHHHHHRRRRRIIIIRRRMRRRIIIIIIIIRRRRRRRRMMLLLLLLLLBBLLLLLLLLLLHHHHRRRRRRRRRRRRRRRRRRRRMMMLLLLLLL
1: LLHHHHHHHHRRRRRMMMMLLLLLLLLLLLLIILBB-L-L-LLL-HHHH-R-R-RRRR-IIIRRRBRRRRRBRRRMMMRRRR-IRMMRRRRRRRREE
2: LLHHHHHHHHRRRRRMMMMLLLLLLLLLLLLIILBBLLLLLLLLLLHHHHRRRRRRRRRRRRIIIRRRBRRRRRBRRRMMMRRRRMMMMMRRRRRRRR-R
```

Supplementary Figure 2: **bpRNA-align results for aligning two 23S structures from the bpRNA-1m meta database. A.** 23S RNA structure with ID bpRNA_CRW_55302. **B.** 23S RNA structure with ID bpRNA_CRW_55267. **C.** Alignment results between the two 23S structures.

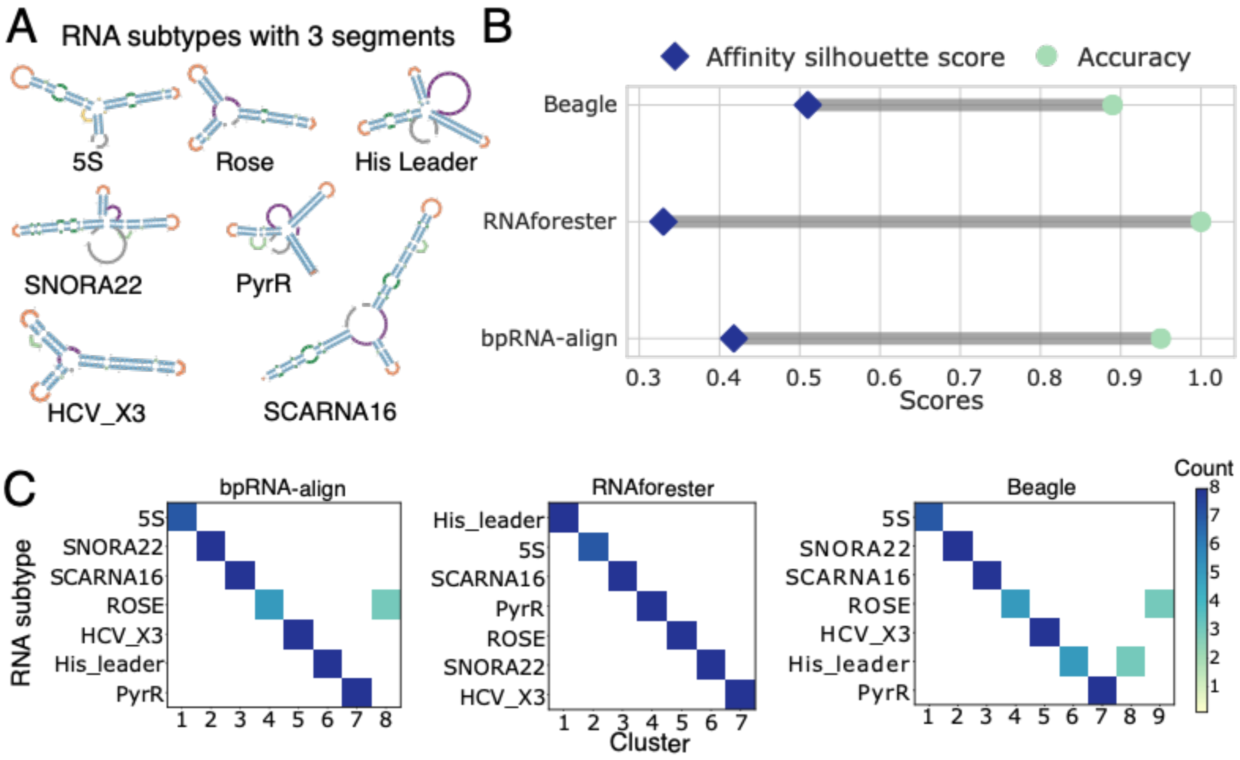


Supplementary Figure 3: **Comparison of structure comparison approaches applied to structures containing 3 segments. A.** RNA subtypes utilized in development of the dataset. **B.** Silhouette and accuracy scores resulting from each structure comparison method. **C.** Confusion matrix results for the methods compared.

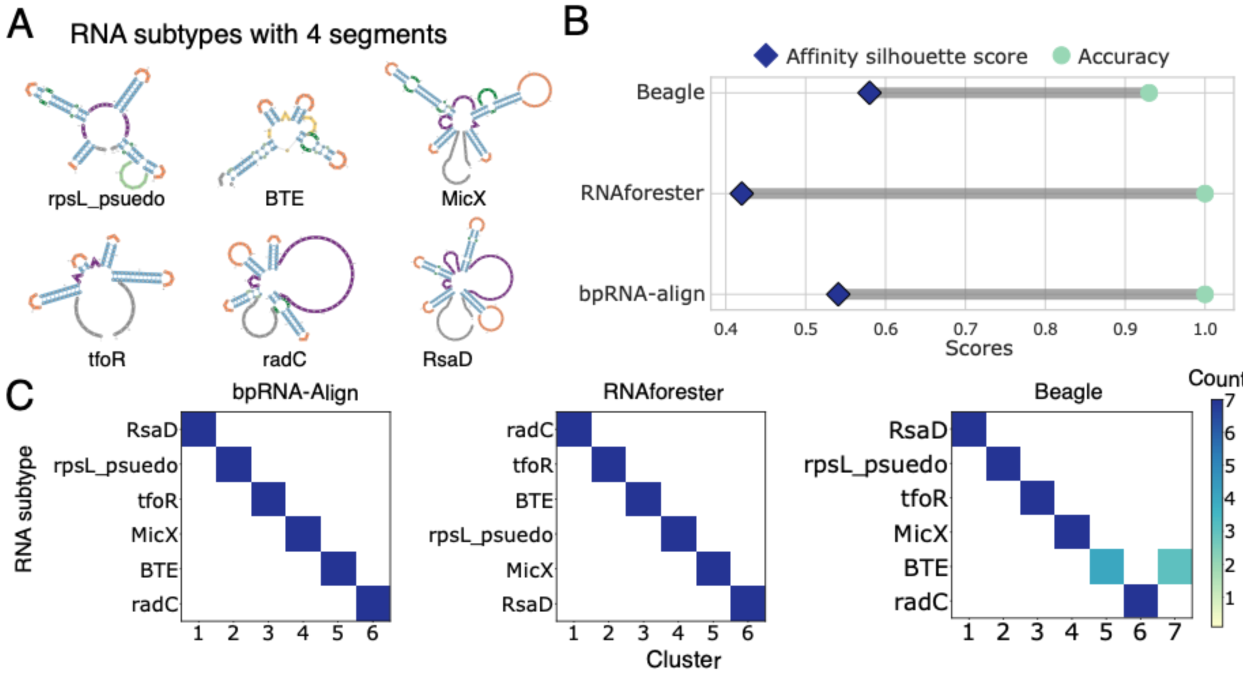


Supplementary Figure 4: **Comparison of structure comparison approaches applied to structures containing 4 segments. A.** RNA subtypes utilized in development of the dataset. **B.**

Silhouette and accuracy scores resulting from each structure comparison method. **C.** Confusion matrix results for the methods compared.

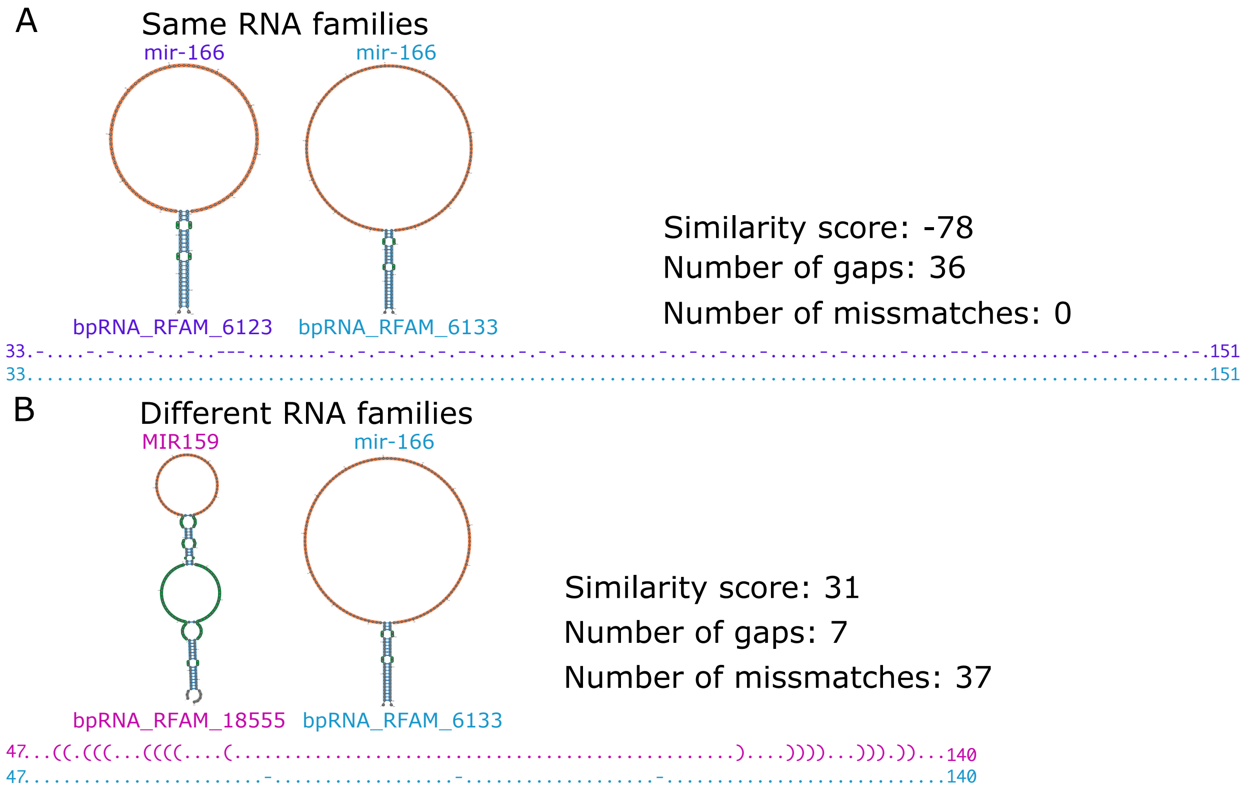


Supplementary Figure 5: **RNAforester alignment examples for structures within the same family, and within different families. A.** RNAforester alignment of mir-166 structures bpRNA_RFAM_6123 and pRNA_RFAM_6133. **B.** RNAforester alignment of structures from two different families (mir-166 and MIR159) with IDs bpRNA_RFAM_6123 and bpRNA_RFAM_6133.
